# Generation and Selection of a Panel of Pan-Filovirus Single-Chain Antibodies using Cell-Free Ribosome Display

**DOI:** 10.1101/327296

**Authors:** Adinarayana Kunamneni, Elizabeth C. Clarke, Chunyan Ye, Steven B. Bradfute, Ravi Durvasula

## Abstract

Filoviruses, which include ebolaviruses and marburgvirus, can cause outbreaks of highly lethal hemorrhagic fever. This disease causes significant morbidity and mortality in humans and non-human primates, with human fatality rates reaching 90% during some outbreaks. Currently, there are a lack of licensed vaccines or antivirals for these viruses. Since early symptoms of filovirus infection mimic more common diseases, there is a strong unmet public health and biodefense need for broad-spectrum filovirus rapid diagnostics. We have generated a panel of mouse single-chain Fv-antibodies (scFvs) to filovirus glycoproteins (GPs) using cell-free ribosome display and determined their cross-reactivity profiles to all known filovirus species. Two scFvs (4-2 and 22-1) were able to detect all known *Ebolavirus* and *Marburgvirus* species. This is the first report on ribosome display scFvs that can detect a broad set of filovirus GPs, which demonstrates their potential use in the development of a new generation of rapid diagnostic immunoassays.

## Author summary

Filoviruses, such as Ebola virus, cause severe and often fatal hemorrhagic fever in humans. The current Ebola virus outbreak is a health crisis with global repercussions and underscores the need for rapid and inexpensive diagnostics to detect filovirus infection. For these reasons, we used a ribosomal display method to rapidly generate single-chain antibodies (scFv’s) for use in diagnostics against all six pathogenic filoviruses from mice immunized with EBOV virus-like particles (VLPs). We have successfully isolated two scFvs that can detect all known *Ebolavirus* and *Marburgvirus* species using this method. This method is inexpensive, rapid, and can be used to quickly develop repertoires of high-affinity antibodies for detection of a broad set of filovirus GPs.

## Introduction

The recent Ebola virus pandemic was a health crisis with global repercussions (PMID 27813879). Rapid spread of the disease within the epidemic regions coupled with migration of infected persons has underscored the need for rapid, robust and inexpensive diagnostic tools. There are six filoviruses known to be pathogenic in humans: four ebolaviruses (Ebola (EBOV), Sudan (SUDV), Tai Forest (TAFV), and Bundibugyo (BDBV) and two marburgviruses (Ravn (RAVV) and Marburg (MARV)) (PMID 21046175). Currently, diagnostics for active infection are largely based on polymerase chain reaction (PCR) platforms. While these are very sensitive and portable units have been made, they are expensive and require considerable training for proper use. There are ELISA-based detection platforms for EBOV, SUDV, and BDBV VP40, but sensitivity and specificity is low relative to PCR-based assays [1]. Furthermore, most of the diagnostic platforms of all types have centered on detection of EBOV. There are fewer readily available monoclonal antibodies for SUDV, BDBV, TAFV, MARV, and RAVV [2], severely limiting development of antibody-based diagnostics. The WHO has recently established ideal guidelines for Ebola virus diagnostics, which include ease of handling, portability, and rapid results (WHO, 2015)[3]. In 2014, the FDA issued 9 emergency use authorizations for EBOV diagnostics. Of these, 8 were PCR-based, and 1 was an antigen ELISA with variably low sensitivity and low specificity [1]. Additional concerns exist regarding filovirus mutants, which may not be detected by standard assays. The recent EBOV Makona strain had mutated so that multiple known antibody epitopes had changed [4]. This would invalidate certain diagnostic platforms that cannot rapidly be modified to detect emerging viral mutants. In addition, in the presence of selective pressure, filoviruses have been shown to readily mutate [5–9]. There is thus a strong unmet public health and biodefense need for broad-spectrum rapid diagnostics and outbreak control.

Ribosome display is a powerful tool for selecting specific antibodies from a large library in cell-free system. It involves the generation of stable antibody-ribosome-mRNA ternary complexes (ARM) followed by panning of the ARM ternary complexes against an immobilized ligand. The mRNA encoding selected target-binding library members is then recovered as DNA from the bound ternary complexes using *in situ* RT-PCR and either analyzed directly by sequencing for *E. coli* expression to screen for desirable candidates or subjected to subsequent rounds of panning for enriching ligand-specific binding molecules. The use of *in situ* RT-PCR also facilitates automation of the entire ribosome display process. Ribosome display has been explored to rapidly produce novel molecules from very large libraries, in combination with next-generation sequencing (NGS) technology. This technology can also be combined with DNA mutagenesis for antibody evolution *in vitro* to produce high-affinity antibodies. More recently ribosome display have been used to obtain ligands with sub-picomolar affinities for the relevant antigen, outperforming the affinities of most conventional mAbs [10]. In addition to antibodies, ribosome display has been explored to select for scaffolds, peptides, ligand-binding molecules, receptors, enzymes, membrane proteins and engineered vaccines [11]. It can also be combined with protein microarrays to allow library-versus-library screening for high-throughput generation of antibodies and genomic discovery of protein-protein interactions [12, 13].

Single chain Fvs (scFvs) are recombinant antibody fragments, which consists of a full variable (epitope-binding) region of an immunoglobulin heavy chain (VH) and the corresponding variable region of an immunoglobulin light chain (VL) variable domains that are connected by a flexible peptide linker [14–16]. They represent the smallest functional antigen-binding domain of an antibody necessary for high-affinity binding of antigen [17].

In this study, we report the use of a rapid, cell-free ribosomal display system to isolate a panel of scFv’s that can bind the glycoprotein (GP) of EBOV, SUDV, RESTV, BDBV, TAFV, and MARV. The resulting antibodies have been characterized in a range of *in vitro* assays which demonstrates their diagnostic potential.

## Results

### Anti-ZEBOVGP scFv antibodies generated from cell-free ribosomal display

To determine whether ribosomal display is suitable for generation of antibodies against filoviruses, we vaccinated mice with virus-like particles (VLPs) consisting of EBOV VP40 and GP. The use of VLPs as a protective vaccine is well-established, as we and others have shown [18, 19].

For ribosome displayed scFv antibody libraries, the immunoglobulin VH and VL regions joined to a 20 amino acid flexible linker [(G4S)4] were constructed using RNA isolated from the spleen of mouse #1 (Figs 1 and 2 and S1 Table). cDNA was synthesized using a single consensus primer MVKR (specific reverse primer) and then pooled and directly used as a template for PCR amplification of the VH and VL chain gene fragments. The gene fragments from mouse were pooled in one reaction tube together with primers that contain overlapping linker sequences which allow the VH and VL gene segments to be assembled by overlap extension to form strep tag-conjugated scFvs (Fig 2). The amplified PCR product was the expected size of about 900 bp. The final DNA template encoding the library flanked by a T7 site was used in an *in vitro* ribosome display with a single selection step with recombinant EBOV GP as outlined in Fig 1.

**Fig 1.**
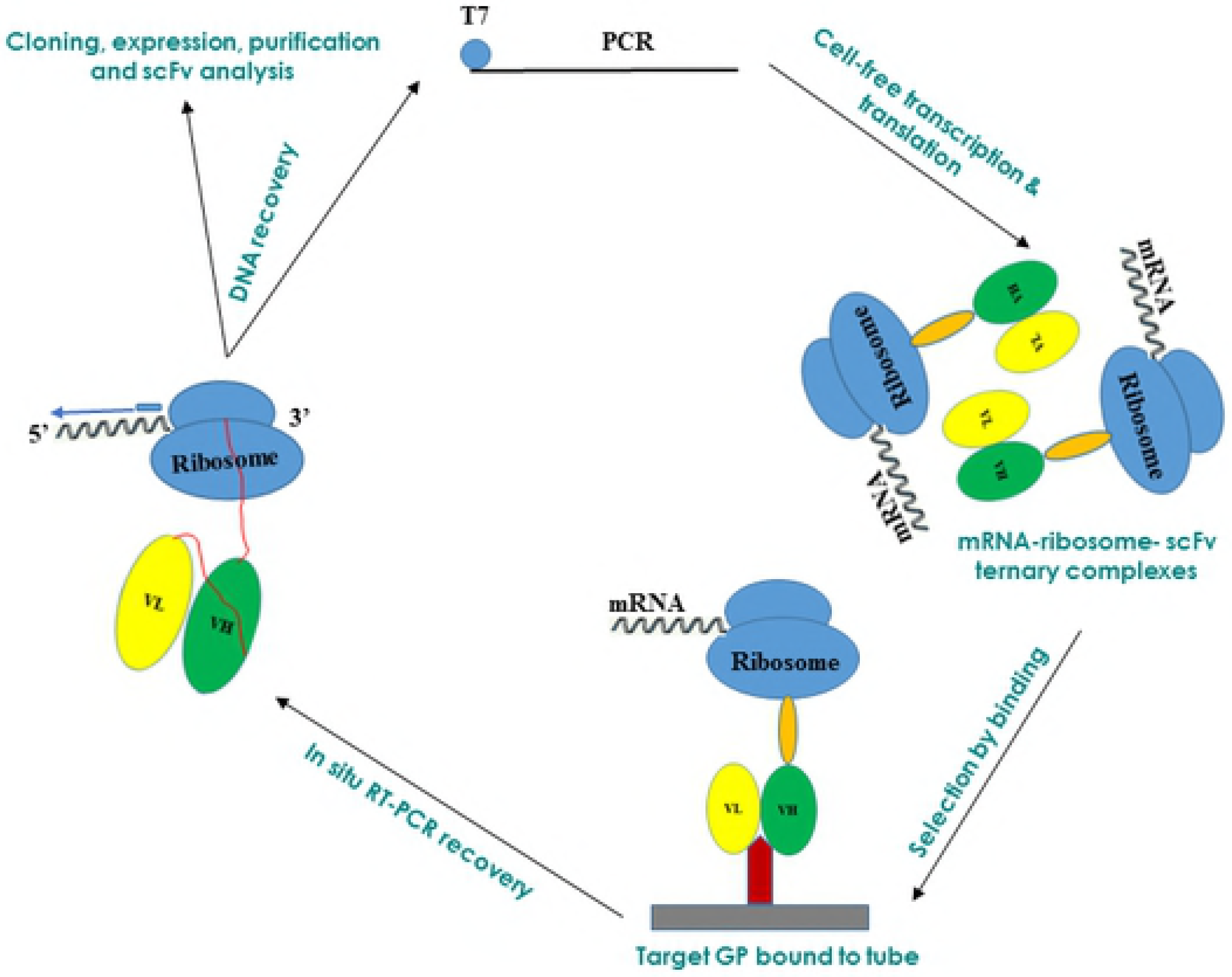
Graphical overview: scFV’s were generated using cell-free ribosomal display from mice immunized with filovirus VLPs.

**Fig 2.**
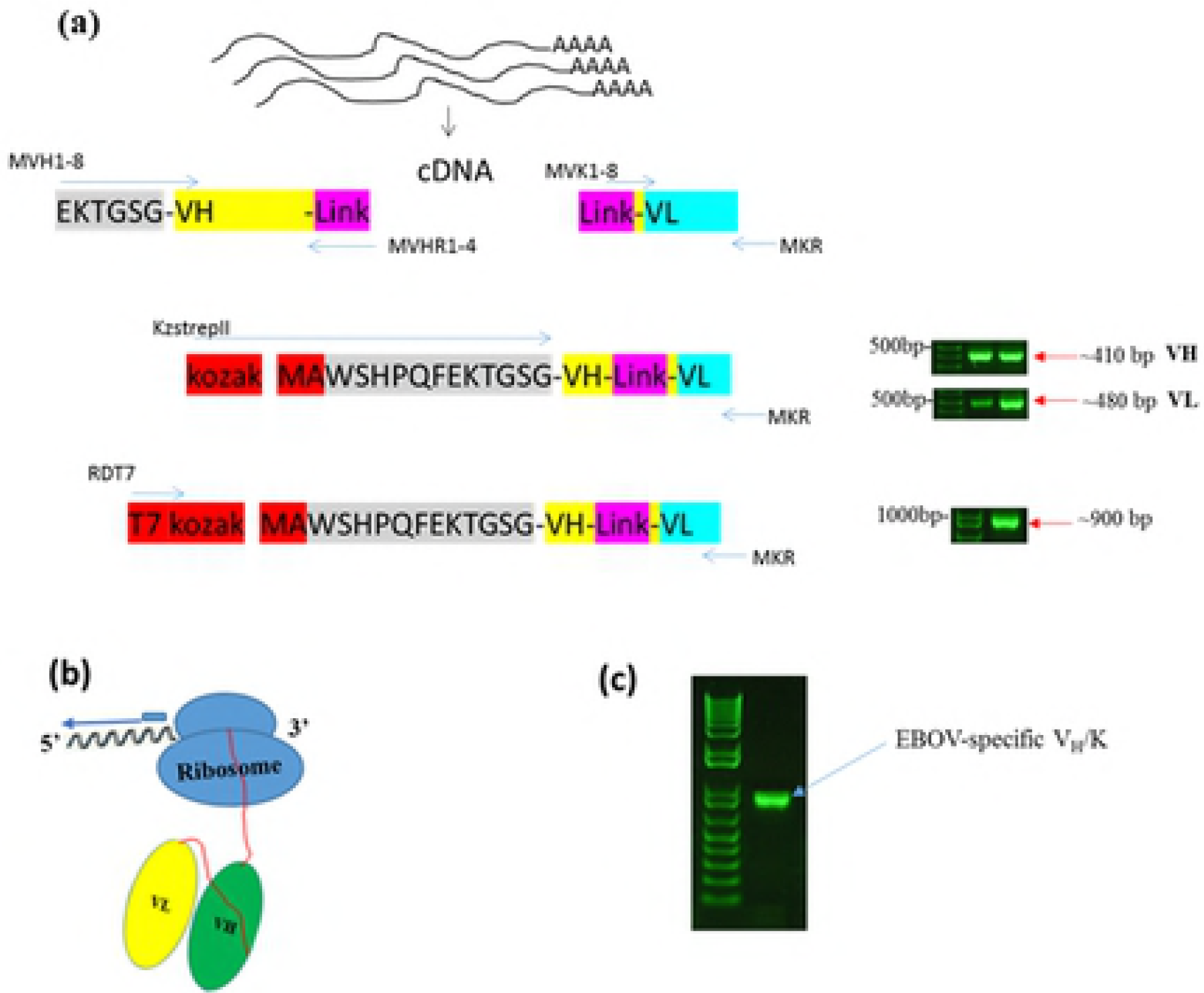
(a) Schematic representation of making scFv library from mouse spleen. (b) Schematic of stalled ARM complex and position of primers used for RT-PCR recovery in the 1^st^ cycle of ribosome display. T7 is the 5’ primer and MKR is the 3’ primer. **(c) Analysis of RT-PCR recovery of V_H_/K cDNA in the 1^st^ cycle.** V_H_/K complexes were bound to EBOV GP coated to PCR tube through 1st cycle of selection and recovery.

After selection, the mRNA was recovered as DNA from the bound ternary complexes by *in situ* RT-PCR (Fig 2 B and C). The PCR product was then cloned into pGEM-T vector DNA from ten randomly chosen clones from the libraries was sequenced and were aligned using Clustal Omega. Their amino acid sequences were deduced and three complementary determining regions (CDRs) and four framework (FW) regions were identified in each of the heavy (VH) and light (VL) chain fragments. A 20 amino acid [(G4S)4] linker was also present. Following alignment with each other, significant diversity in the VH and VL chain was observed especially in the CDRs. Variability was also noted in the framework regions. No two clones had identical VH or VL fragments. The aligned amino acid sequences of ten clones from library are shown in S1 Fig. The framework regions (FRs) and CDRs were determined by the IMGT information system (http://imgt.cines.fr/)(IMGT®/V-Quest)[20]. The length of CDRs was variable. The length of CDR1 VH, with an average length of 8 amino acid residues, CDR2VH with an average length of 8 residues, CDR3 VH ranged from 10 to 18 amino acid residues, with an average length of 13 residues, CDR1 VL with an average length of 9 residues, while 3 amino acid residues were found in CDR2VL and 9 amino acid residues were found in CDR3 VL.

After expression of scFv clones, samples were harvested, lysed, purified, and fractions were analyzed by SDS-PAGE (to confirm the integrity and purity) followed by Western blot. A prominent band of about 33 kDa was confirmed the identity of the purified protein on Western blot (Fig 3A).

**Fig 3.**
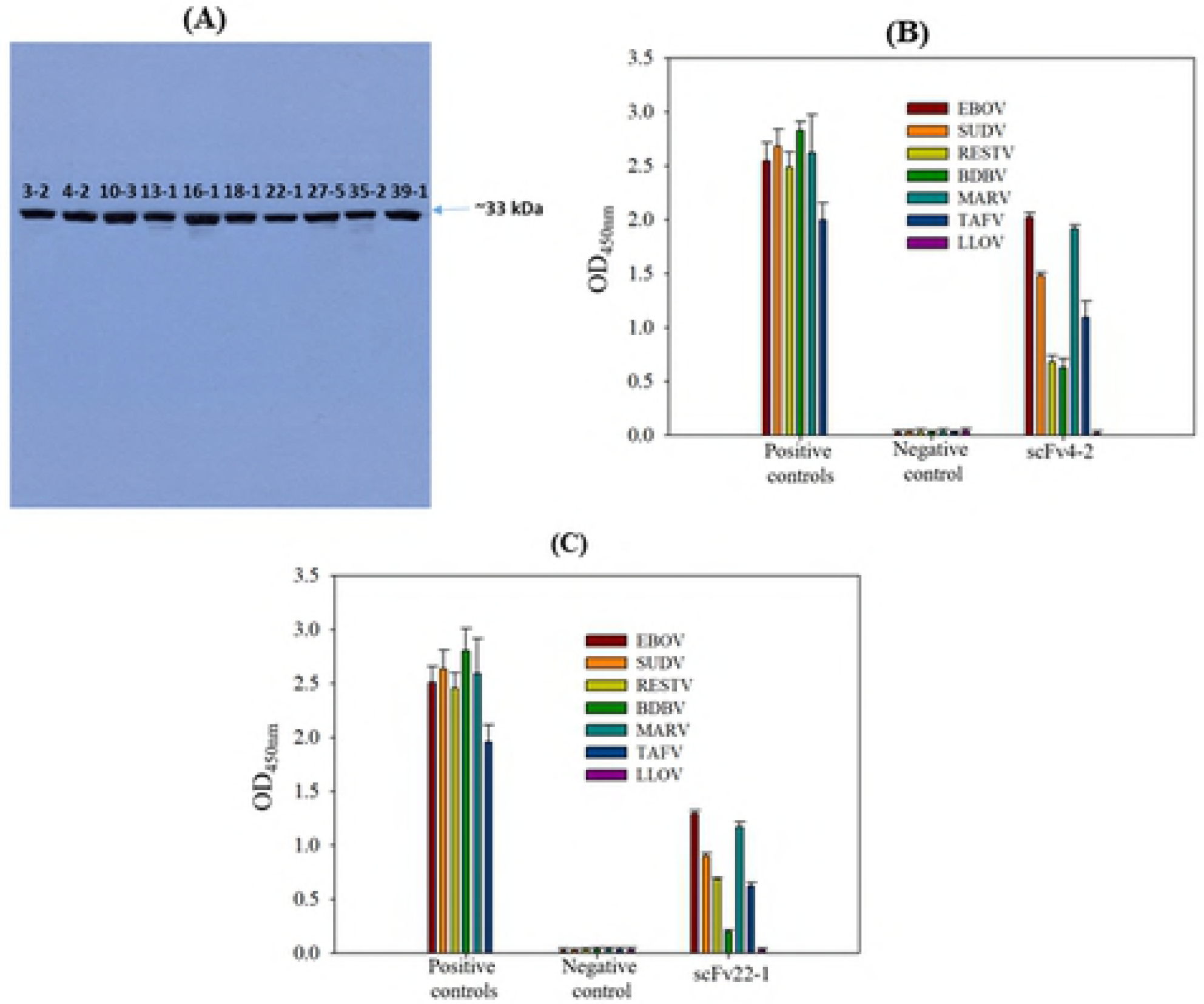
(A) Western blot of purified scFvs (3-2, 4-2, 10-3, 13-1, 16-1, 18-1 22-1, 27-5, 35-2 and 39-1). ELISA results for purified novel anti-EBOV scFv4-2 (B) and scFv22-1 (C) (generated by ribosome display), commercial monoclonal (Mouse anti-EBOV antibody 6D8, and Mouse anti-SUDV ebola virus antibody) and polyclonal (Rabbit anti-RESTV GP, Rabbit anti-BDBV GP, Rabbit anti-MARV GP, and Rabbit anti-TAFV GP) antibodies as positive controls, and an scFv negative control (Mouse anti-PilB scFv21 antibody).

### Specificity, cross-reactivity and affinity of GP-specific scFvs

The scFvs were screened by ELISA for their specificity and cross-reactivity with recombinant GPs of the other known filoviruses in the genus Ebolavirus (SUDV, TAFV, BDBV and RESTV) and Marburg virus (MARV). Several different profiles for the cross-reactivities of these antibodies were found (S2 Table and Fig 3B and C). Two scFvs (EBOVGP4-2 and EBOVGP22-1) reacted strongly with all tested GPs of Ebolavirus and Marburg species. Six scFvs (EBOVGP3-2, EBOVGP13-1, EBOVGP16-1, EBOVGP18-1, EBOVGP35-2, and EBOVGP35-1) bound weakly to GPs of some viruses in addition to EBOV, EBOVGP scFv3-2 reacted only to TAFV and MARV, and EBOVGP scFv27-5 didn’t react with any GPs. Importantly, these different reactivity profiles enabled us to distinguish the known Ebolavirus species by using scFvs (4-2 and 22-1), as shown in Fig 3B and C and S2 Table. Representative scFvs for each obtained cross-reactivity profile showing the highest OD values were selected for dose response relationship studies.

Dose response curves were determined for scFvs EBOVGP4-2 and EBOVGP22-1 (Fig 4A&B), (S3-S6 Tables, highlighted rows) Both scFvs bound tightly to all six GPs, with EC_50_s ranging from 4 to 12 μg/mL. EBOVGP scFv4-2 and EBOVGP scFv22-1 showed the strongest binding to all six ebolavirus species, with EC_50_s below 9 and 12 μg/mL, respectively. The EBOVGP-scFv 4-2 demonstrated about almost 2-fold enhanced binding compared to EBOVGP-scFv 22-1. Binding constant (KD) determinations were estimated by fitting a dose response curve and the KD estimates for the scFvs against all six GPs are shown in S7 Table. The highest affinity scFv was the anti-EBOVGP scFv4-2, which was about 2-fold greater towards all six GPs than the anti-EBOVGP scFv22-1.

**Fig 4.**
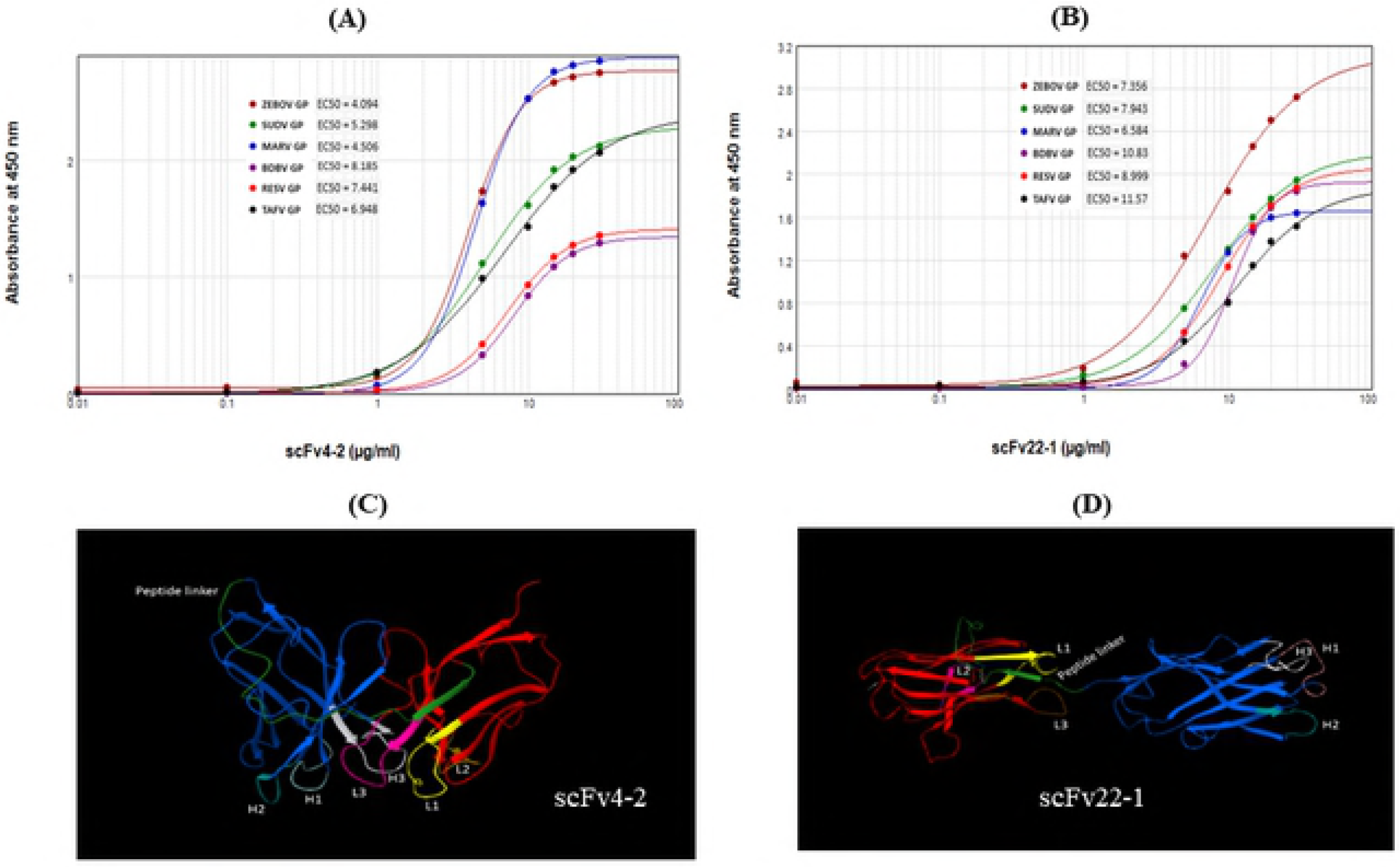
(A&B) Reactivity of mouse ZEBOV GP scFvs (4-2 and 22-1) to filovirus glycoproteins and dose response binding of the indicated antibodies to GPs of EBOV, SUDV, MARV, BDBV, RESTV, and TAFV. Values are optical density at 450 nm (OD450) values from three ELISA experiments performed over the indicated range of antibody concentrations. The EC50s (in micrograms per milliliter) for binding of each antibody to the respective antigen are shown in each panel. Predicted Ribbon representation (C & D) of anti-ZEBOV scFv antibody molecules (4-2 & 22-1) in 3D mode.

To assess the ability of the anti-EBOVGP scFvs to recognize the six filovirus GPs in a denatured format, western blotting was performed on purified GPs. Anti-EBOVGP scFvs 4-2 and 22-1 binding to ebolavirus GPs was completely lost upon denaturation, while binding to GP_MARV_ was not affected (Fig 5). The reactivities of these GP-specific scFvs were further tested by immunoprecipitation analysis (IP) using lysates of filovirus GP expressed in 293T cells. The results demonstrated that the anti-EBOVGP scFvs were able to immunoprecipitate filovirus GPs, confirming the ELISA data. (Fig 6).

**Fig 5.**
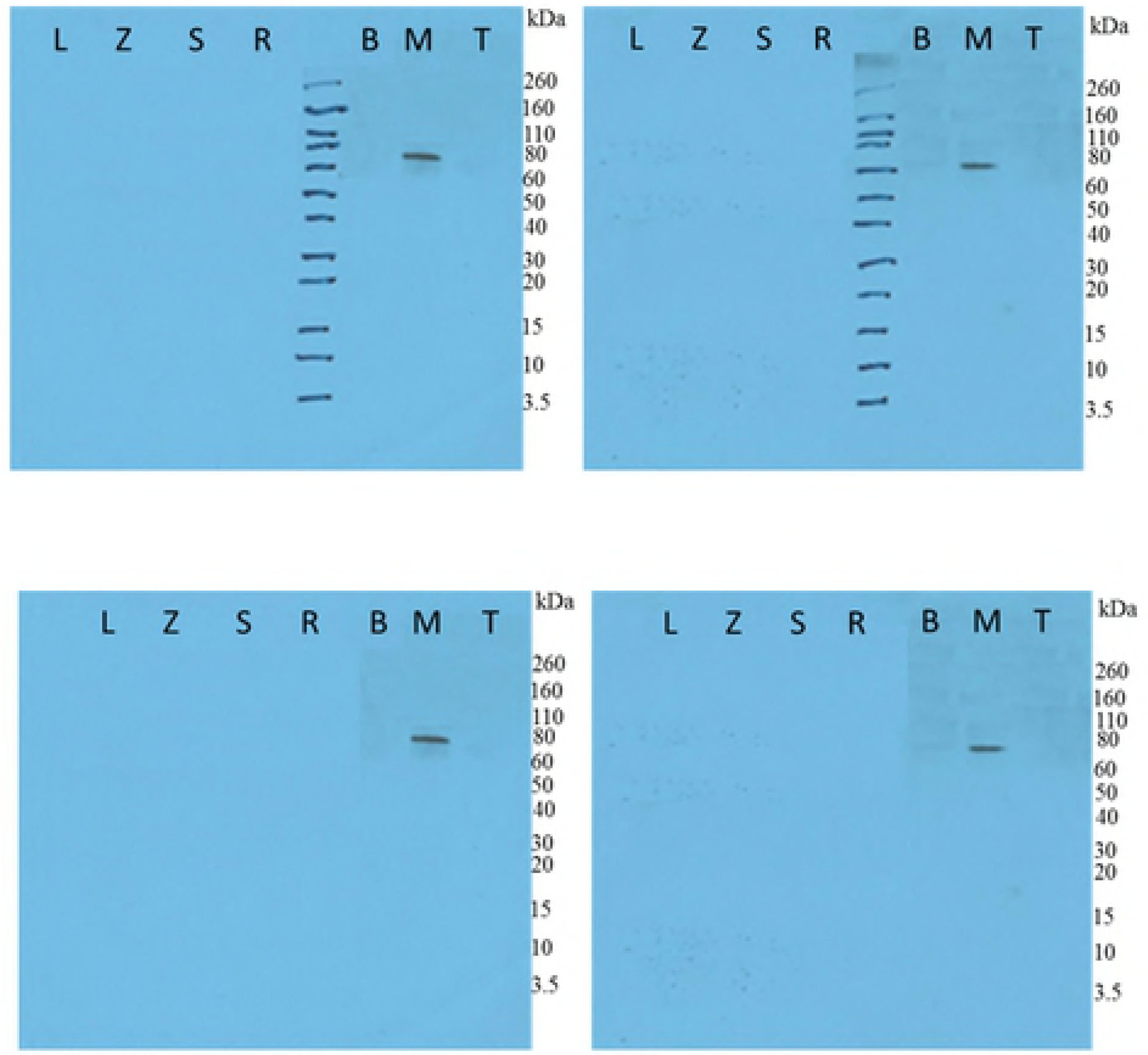
Denaturation eliminates binding of scFV4-2 and scFv 22-1 for the ebolavirus GPs. The purified proteins of LLOV, (L) EBOV (Z), SUDV (S), RESTV (R), BDBV (B), MARV (M) and TAFV (T) GPs without TM were probed with soluble scFvs 4-2 and 22-1.

**Fig 6.**
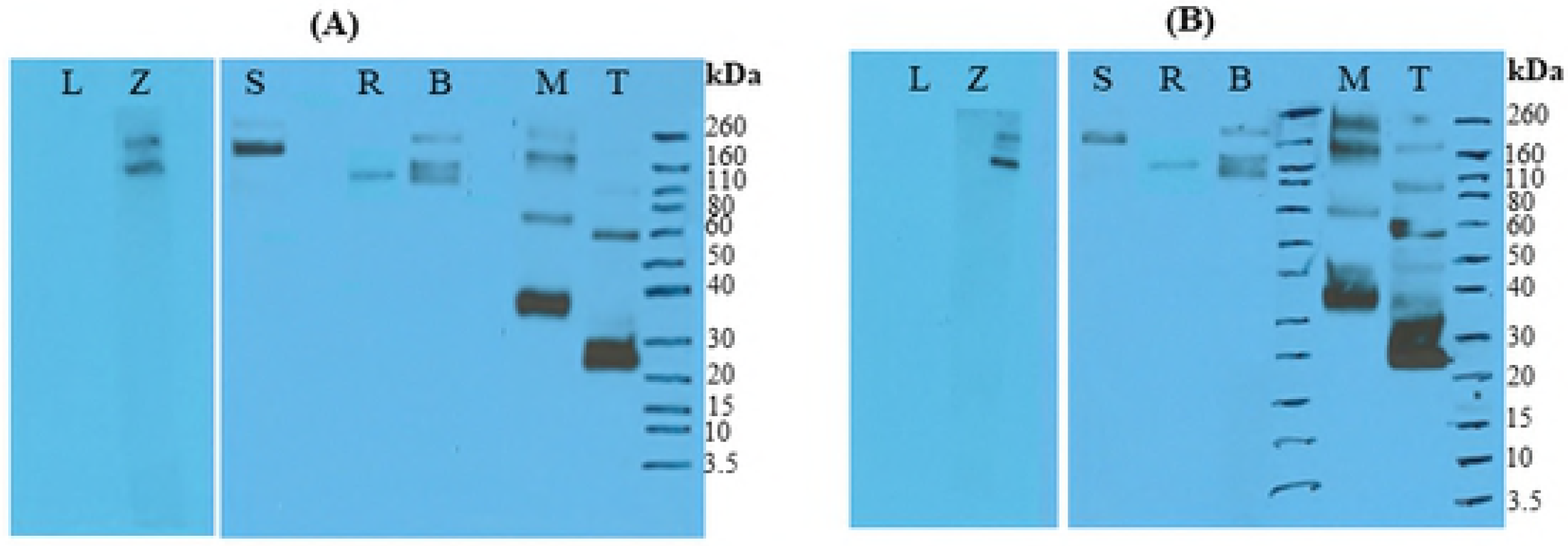
Immunoprecipitation of filovirus GPs by scFvs 4-2 and 22-1. The lysates of LLOV, (L) EBOV (Z), SUDV (S), RESTV (R), BDBV (B), MARV (M) and TAFV (T)-infected 293T cells were immunoprecipitated with soluble scFvs 4-2 and 22-1 followed by SDS-PAGE analysis and western blot.

An immunofluorescence assay (IFA) was performed to confirm the recognition of scFvs with the proteins of GP_EBOV_, GP_SUDV_, and GP_MARV_ in its natural membrane associated trimeric structure with 293T cells transfected with the GP_EBOV_, GP_SUDV_, and GP_MARV_ expressing plasmids (Fig 7). The results demonstrated that these two scFvs (4-2 and 22-1) effectively recognized and stained the GP_EBOV_, GP_SUDV_, and GP_MARV_ expressed in the 293T cells. The staining ability of scFv4-2 was superior to scFv22-1 and these are comparable to the results of positive controls in the IFA.

**Fig 7.**
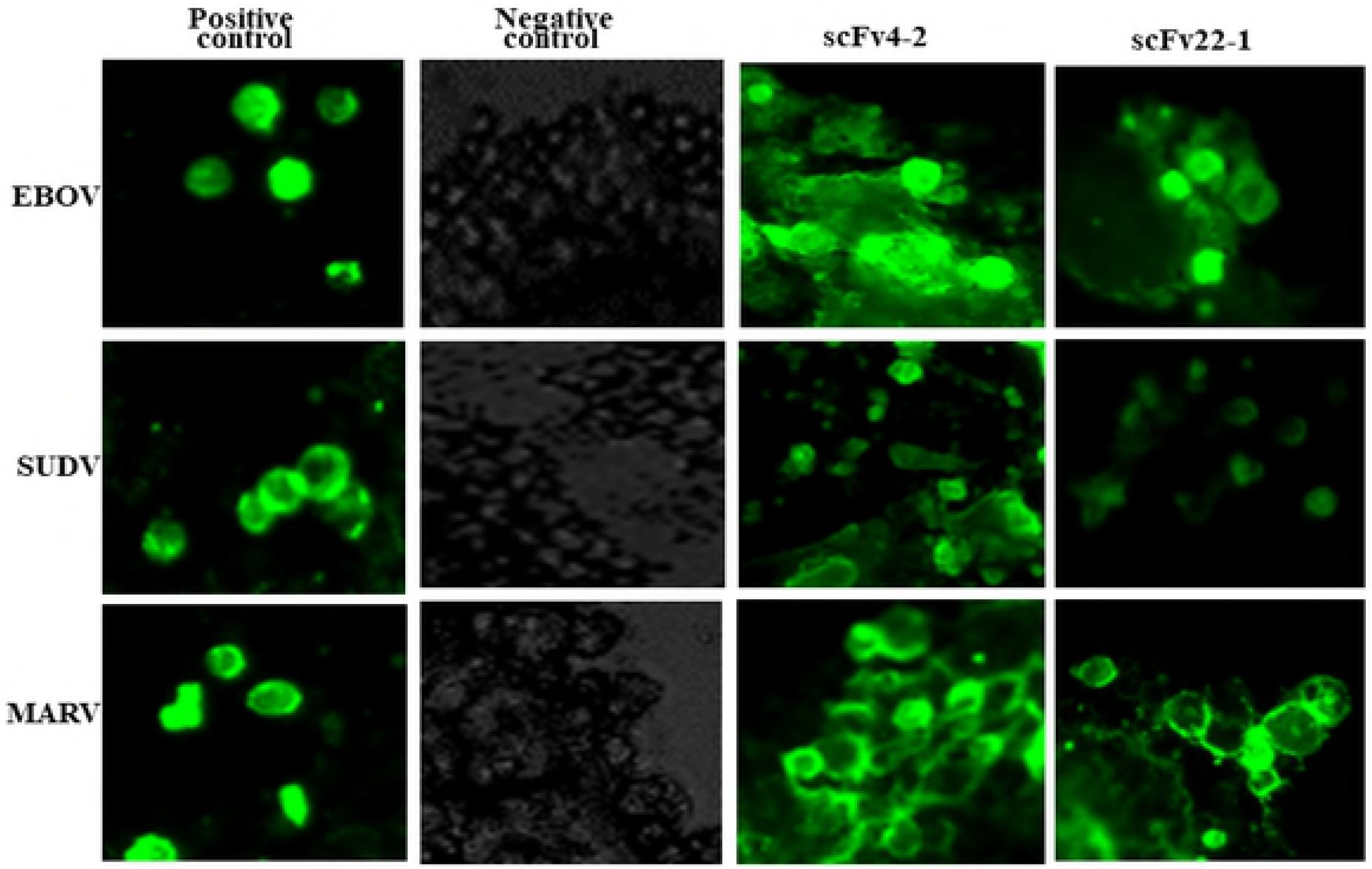
The scFvs 4-2 and 22-1, and positive controls recognize the natural forms of GP1,2_ZEBOV_, GP1,2_SUDV_, and GP1,2_MARV_ by IFA.

## Discussion

The latest Ebola virus disease (EVD) outbreak in West Africa has highlighted the urgent need for rapid and effective diagnostics against filoviruses. Here we report novel murine scFv antibodies with a range of cross-reactivity to five species of ebolavirus and to marburgvirus. Two of these scFvs can cross-react to all six known pathogenic filoviruses, representing pan-filovirus binding. This is the first report of the use of cell-free ribosome display and epitope mapping to generate broadly-reactive scFvs against filoviruses.

Of the 10 anti-EBOV GP scFvs tested by ELISA, there were two that were highly cross-reactive to all ebola- and marburgvirus GPs (S2 Table and Fig. 3B and C). Holtsberg et al. [21] demonstrated that a single MAb (m21D10) reactive to three GP (EBOV, SUDV, and MARV) molecules and six clones reactive to both EBOV and SUDV but nonreactive to MARV were identified. Keck et al. [22] identified a set of broadly reactive pan-ebolavirus MAbs with strong binding to EBOV, SUDV, BDBV, and RESTV GP; two MAbs also exhibited weak binding to MARV GP. Other studies have also found broadly cross-reactive ebolavirus GP antibody (PMID 28525756 and 28525755). Our binding data showed that the cross-reactive scFvs, 4-2 and 22-1, exhibited moderate to higher binding with submicromolar EC_50_s to five ebolavirus and a marburgvirus GPs (Fig. 4A and B). Our results are comparable to the results from the dose-dependent binding of mAb m21D10 to EBOV, SUDV, and MARV GP [21] and FVM04, exhibited balanced binding at subnanomolar concentrations to four ebolavirus species tested, shows weak binding to MARV [22]. scFv4-2 antibody showed an approximate 2-fold high affinity to all GPs in comparison to the scFv22-1 antibody (S7 Table and Fig. 7). Rodríguez-Martínez et al. [23] reported that the scFv-13C6 mAb exhibited the highest EBOV GP-binding activity, followed by scFv-13F6 and Fab-KZ52 in ELISA experiments.

Western blot suggests that the epitope of these scFvs is completely lost in denaturation (Fig. 5). Interestingly, these scFvs recognized MARV, suggesting that these scFvs bind to a different epitope (data not shown). These scFvs bound to all six GPs in an ELISA, but not a Western Blot (except for MARV GP), suggesting these can recognize GP in its natural conformation and binds to a conformational epitope.

On IP analysis using lysates of filovirus GPs produced in 293T cells (Fig 6), the cross-reactivity profiles and virus specificities were similar to those obtained by ELISA. These scFvs, cross reactive to all known ebolaviruses, may be very important for the development of Ebola and Marburg virus antigen detection assays.

Although a previous study demonstrated in vivo efficacy of anti-marburgvirus GP scFvs [24], this is the first report of a scFv antibody (4-2 or 22-1) that exhibits broadly cross-reactive activity against filovirus GP. These scFvs also have the potential for identification of as yet unknown Ebolavirus species. During the first reported outbreak of BDBV, initial diagnosis was made using an antigen capture ELISA using a combination of anti-ZEBOV, SUDV and RESTV mAbs as well as hyper-immune polyclonal anti-EBOV rabbit serum, whereas the specimens were initially negative when tested by RT-PCR [25]. This highlights the value of scFvs to potentially identify new Ebola and Marburg virus species. The scFvs generated in this study may therefore offer some advantage in generating a highly sensitive assay for diagnostic use.

In summary, a panel of EBOVGP-specific scFvs from ribosome display and their cross-reactivity profiles to all known human pathogenic filoviruses was established. This is the first report on ribosome display scFvs that can detect all known ebolaviruses. These broadly cross-reactive scFvs produced in this study have potential use in the development of new generation rapid diagnostic tests for the detection of filoviruses.

## Materials and Methods

### Ethics Statement

Mice experimental procedures were approved by the Institutional Animal Care and Use Committee (IACUC) at the University of New Mexico. Animal research performed at the University of New Mexico was conducted under an IACUC approved animal protocol (protocol #15-200336-HSC)in compliance with the Animal Welfare Act and other federal statutes and regulations relating to animals and experiments involving animals and adheres to principles stated in the *Guide for the Care and Use of Laboratory Animals* [26]. The University of New Mexico is fully accredited by the Association for the Assessment and Accreditation of Laboratory Animal Care International.

**Plasmids, Strains and Reagents.** All reagents used in the study were commercially available and were of reagent grade or better. All restriction enzymes and DNA modification enzymes were of molecular biology grade. All primers were purchased from Invitrogen. pGEM-T easy cloning vector and TNT T7 Quick for PCR DNA kit (rabbit reticulocyte cell free extract) were purchased from Promega. Plasmid pBAK1, previously constructed in our laboratory is based on pET-26b vector (Novagen). Rabbit anti-Strep tag II Polyclonal Antibody Affinity Purified HRP conjugated, mouse anti-his antibody and rabbit anti-mouse AP antibody were purchased from GenScript Inc., Invitrogen and Sigma-Aldrich, respectively. AP detection reagent kit was purchased from Novagen (USA).

### Cell lines and plasmids

HEK293T cells (American Type Culture Collection) were maintained in Dulbecco’s Modified Eagle Medium (DMEM; Gibco) + 10% fetal bovine serum (FBS) + 1% penicillin/streptomycin. Filovirus GPs were encoded in pcDNA3.1 plasmids.

### Ebola VLPs

eVLPs were prepared essentially as previously described, with minor modifications [19, 28]. Plasmids expressing EBOV-Yambuku GP1,2 and VP40 were transfected into 293T cells using JetPrime, according to the manufacturer’s protocol. Three days after transfection, supernatants were harvested and centrifuged at 9,000 g for 2 hours. VLP pellets were resuspended in PBS and layered for a discontinuous 60/30/10 gradient, and ultracentrifuged for 18 hours at 150,000 g. Lower bands were collected, diluted in PBS, and centrifuged for 2 hours at 100,000 g. Pellets were resuspended in PBS, and quantitated.

### Immunization of mice

C57BL/6 mice were obtained from The Jackson Laboratory. Mice were vaccinated i.p. with 10 ug of EBOV virus-like particles (VLPs) mixed with alum twice at 3-wk intervals. Splenocytes of individual mice were placed in 10 mL of TRIzol for use in RNA isolation approximately 6 weeks after the last vaccination.

### Generation of single chain antibodies by cell-free ribosome display

#### Antibody library construction

Total spleen RNA was prepared as described by Andris-Widhopf *et al.* (2001). Mice spleens were minced and homogenised in 10 mL TRIzol (Invitrogen). The total RNA pellet was air-dried and resuspended in 500 μL of nuclease-free water (stored at −80°C). Complementary DNA (cDNA) was synthesized from approximately 25 μg of total RNA using a SuperScript II RT (Invitrogen) following the manufacture’s instructions provided.

For antibody library construction, the PCR primers were based on published sequences [29] with minor modifications (see S1 Table). The primers were designed to introduce in-frame NcoI and NotI restriction sites to the 5’ end of the VH sequence and to the 3’ end of the VL sequence, respectively. The VH_F/VH_R and VL_F/VL_R sets of primers (see S1 Table) were used for PCR amplification of VH and VL gene segments using the cDNA template. The VH_R and VL_F set of primers (see S1 Table) were used to introduce overlapping sequences which enabled the scFv gene fragments to be assembled by overlap extension PCR and these primers encode a 20 amino acid linker sequence (G4S)4. The amplified heavy and light-chain products were purified and pooled, and an aliquot of light and heavy-chain templates was subjected to overlap extension PCR amplification using Link to introduce Strep II tag and Kozak sequence on the 5’ end and an overlap extension on the 3’ end to facilitate joining to the variable heavy-chain libraries using MKR and KzSTREPII. Finally, the PCR product encoding all the variable heavy-chain and light-chain combinations was amplified with primers RDT7 and MKR to introduce T7 site into Strep tag II-conjugated VH-VL library and to produce the DNA encoding the anti-Ebola immunoglobulin scFv libraries. The initial PCR amplification reactions were performed at a 52°C annealing temperature with 30 cycles, and the subsequent library assembly step used 16 cycles with *GoTaq* DNA polymerase and 20 pmoL of each primer pair per reaction. DNA fragments were resolved by gel electrophoresis on 1% (wt/vol) agarose gels. DNA isolation from agarose gels was carried out following QIAquick DNA gel purification kit instructions. The final purified PCR product ~0.9 kb is a template for Ribosome Display.

#### Cell-free Ribosome display technology

To select specific antibody fragments, we have used a modified eukaryotic ribosome display with slight modifications[30]. *In vitro* transcription and translation reaction was based on a coupled rabbit reticulocyte lysate system (Promega’s TNT quick-coupled transcription-translation system) and performed according to the supplier’s protocol. The PCR-generated DNA library of antibody-coding genes derived from mouse 1 were expressed in this lysate system. Briefly, 50 μL of transcription/translation mixture containing 40 μL of TNT T7 Quick Master Mix, 2 μL of DNA library (0.1 to 1.0 μg), 1 μL (1 mM) of methionine, 1 μL of DNA enhancer, and 6 μL of water were added, and the reaction mixture was incubated at 30°C for 90 min. Then, 5 μlLof RNase-free DNase I (Roche) (2,000 U/ml) was added, and the mixture was incubated for 20 min at 30°C (in order to remove the DNA template so that subsequent PCR only picks up pulled down RNA sequences).

To select specific antibody fragments the 0.5 mL PCR tubes were coated with 1 μg/mL of the recombinant Zaire Ebov GP in 100 μL PBS at 4 °C overnight. Protein coated tubes were washed with PBS and blocked with 100 μL of molecular biology grade Bovine Serum Albumin (BSA) in PBS (10 mg/mL) (New England Biolabs) for 1 hour at room temperature. The translation/transcription mixture (containing the protein-ribosome-mRNA (PRM) complexes) was added to the washed and blocked protein-coated tubes and incubated on ice for 1 hour. The PCR tubes were washed three times with ribosome display washing buffer (PBS containing 0.01% Tween 20, 5 mM Magnesium acetate and 0.1% BSA, pH 7.4) and two times quick wash with ice-cold RNase-free water, and the retained RNA (antibody sequences) subjected to the following recovery process; *in situ* Single-Primer RT-PCR Recovery was performed in the PCR tubes carrying selected ARM complexes using a SuperScript II reverse transcriptase (Invitrogen, USA).

The obtained cDNA was amplified in a 25 μL PCR mixture for 35 cycles of 30 s at 94°C, 30 s at 52°C, and 1 min at 72°C, and 10 min at 72°C with MKR and KzStrep II using *GoTaq* DNA polymerase (Promega, USA). The RT-PCR product from a single round of ribosome display was purified by agarose gel electrophoresis.

#### Cloning, expression and purification of an anti-Ebov GP scFvs

The RT-PCR product from a third round of ribosome display was cloned into the pGEM®-T Easy vector (T-Cloning® Kit, Promega) according to the manufacturer’s instructions. The ligation products were transformed in pGEM®-T Easy vector *E. coli* cells and positive colonies were chosen by blue-white selection and confirmed by DNA sequencing using T7 and SP6 standard primers (S1 Table). Based on the sequencing results, the clones in right reading frame without stop codon were chosen for prokaryotic expression. EbovGP-scFv is expressed in the form of C-terminal 6xHis fusion from the prokaryotic expression vector pBAK1, previously constructed in our laboratory is based on pET-26b vector (Novagen). This vector has a strong T7 promoter, and is designed to work with Rosetta-gami (DE3) strains of *E.coli.* The protein is induced and expressed at room temperature (25°C), to increase the portion of the correctly folded, soluble EbovGP-scFv.

For the construct of scFvs, DNA was amplified with the forward primer, KzStrepII 5‘CGAATTCCA**<u>CCATGG</u>**CCTGGAGCCATCCGCAGTTCGAGAAGACCGGCAGCGG3’ (with an NcoI restriction sequence highlighted in bold) and the reverse primer, MKR_NotI 5’AGT**<u>GCGGCCGC</u>**AGAACACTCATCCTGTTGAAGCTCTTGACAATGGGTGAAGTTG3’ (with an NotI restriction sequence highlighted in bold). Both set of primers allow these two amplicons (S2 Fig) to be subcloned into a pBAK1-His vector [31] and were transformed into Rosetta gami B(DE3) *E. coli* strain and plated onto LB agar plate with 100 μg/mL of carbenicillin, and grown at 37 °C for 16 hours. The molecular weight and isoelectric points was predicted using ExPASy bioinformatics resource portal (http://web.expasy.org/compute_pi/)[32]. Next day, five colonies were inoculated in 3 mL of LB media with the same antibiotics and grown at 37°C with 225 rpm shaking for 16-18 hours. and the positive clones were selected by restriction analysis with NcoI and NotI and confirmed by DNA sequencing. The 3 mL overnight culture was used to prepare a glycerol bacterial stock and inoculated into 200 mL of LB medium containing the same antibiotics and grown at 37°C with shaking at 225 rpm until OD_600_ reached between 0.4-0.6. The culture was briefly chilled on ice to 25°C and the cells were induced by the addition of IPTG (final concentration 1 mM) and were incubated for 12 hours at 25°C with shaking. Cells were harvested in 4×50 mL tubes by centrifugation at 4000 *g*, 4°C for 20 minutes. Cell pellets were frozen and stored at −80°C before undergoing further processing.

The 50 mL cell pellet from 200 mL culture was re-suspended in 3 mL of lysis buffer (20 mM Tris-HCl, 500 mM NaCl, 20 mM Imidazole, 0.1 % Triton X-100 pH 8.0). Cells were lysed by sonication on ice (6×30 seconds), and were centrifuged at 14,000 *g* for 15 minutes to remove cellular debris. The soluble fraction was filtered through 0.2μm filters and applied to HisTrap excel 1mL column (GE Life Sciences). The column was equilibrated and washed with 20 mM Tris-HCl, 500 mM NaCl, 20 mM Imidazole pH 8.0 and the sample was eluted 20 mM Tris-HCl, 500 mM NaCl, 500 mM Imidazole, pH 8.0 in one elution step. The purification was carried out at a constant flow of 1 mL/min. Fractions of 1 mL were collected through the elution step. The purified protein fractions were kept at 4 °C.

#### Western blot of scFv fragments

The proteins were separated by SDS-PAGE and were blotted onto polyvinylidene difluoride (PVDF) membrane using a iBlot2 Gel TransferDevice (Life Technologies) for 7 min. The membranes were blocked in 5% w/v skimmed milk powder/TBS buffer at room temperature for 1 hour, then incubated with mouse anti-His antibody (GenScript, Piscataway, NJ, USA) at a dilution of 1:5000 for 1 h. After three washes with TBST buffer, HRP-conjugated Rabbit anti-strep tag II antibody (diluted 3/10,000 in TBST) (GenScript, Piscataway, NJ, USA) at room temperature for 1 hour, followed by washing with TBS-T buffer 3x for 10 minutes each. The protein was detected and visualized with ECL detection reagent (Amersham, USA) following the manufacturer’s recommendation.

#### Transfections

All transfections were performed in HEK293T cells seeded in a 6-well plate (PMID 29118454 Cells were transfected with filovirusGPs (sequences listed in PMID 29118454) using JetPrime according to the manufacturer’s recommendations. Cells were stained with antibodies 48 hours after transfection.

### Analysis of purified scFvs

#### Specificity, cross-reactivity and affinity of GP-specific scFvs

To determine binding and cross-reactivity and half maximal effective concentrations, or EC50s, scFvs were tested for binding to the GP antigens (1 μg/ml) of EBOV, SUDV, BDBV and RESTV, as well as to MARV Ravn GP at a concentration range of 30 μg/ml to 0.1μg/ml. ELISAs were performed as follows: Nunc Polysorp ELISA plates were coated overnight at 4°C either with target filovirus GP antigens, blocked with 2% BSA at 37°C for 1 h and incubated with anti-EBOV scFvs in 1% BSA. Commercial monoclonal (Mouse anti-EBOV antibody 6D8, and Mouse anti-SUDV ebola virus antibody) and polyclonal (Rabbit anti-RESTV GP, Rabbit anti-BDBV GP, Rabbit anti-MARV GP, and Rabbit anti-TAFV GP) antibodies were served as positive controls, whereas Mouse anti-PilB scFv21 antibody (unpublished data) as negative control. After 1 hour incubation at 37°C, plates were washed as described above and scFv was detected with 6:10 000 dilution of HRP-conjugated Rabbit anti-strep tag II polyclonal antibody in PBS containing 1% BSA and incubated at 37°C for 1 hour. After washing and tap drying, 100 μL of TMB substrate was added to each well and the plate was allowed to incubate for 15-30 minutes at 25°C. The reaction was stopped with 0.18M H_2_SO_4_, and absorbance reading was measured at 450 nm and these values were transformed using Softmax Pro (7.1) 4 parameter curve-fit (Molecular Devices Corp., CA, USA).

#### Western blot analysis

Different purified pan-ebola and pan-filovirus antigens (LLOV, ZEBOV, SUDV, RESV, BDBV, MARV and TAFV GPs) of 1 μg/mL were separated on 4-12% SDSPAGE and transferred onto PVDF transfer membrane using a iBlot2 Gel TransferDevice (Life Technologies) for 7 min. The membranes were blocked with 5% skim milk in TBS, 0.1% Tween-20 (TBST) for 2 hours followed by incubation with the scFvs 4-2 and 22-1 (1 μg/ml concentration) diluted in TBST for 1 hour. After three washes with TBST, the membranes were incubated with HRP-conjugated Rabbit anti-strep tag II antibody (diluted 3/10,000 in 5 % Milk/TBS) (GenScript, Piscataway, NJ, USA) at room temperature for 1 hour, followed by washing with TBS-T buffer 3x for 10 minutes each. The protein was detected and visualized with ECL detection reagent as described above.

#### Immunoprecipitation assay (IP)

EBOVHEK293, SUDVHEK293, RESVHEK293, BDBVHEK293, MARVHEK293, LLOVHEK293 and HEK293 cell lysates (containing 10 mg protein) were obtained by lysis with NP40 buffer, protease inhibitor cocktail, and PMSF and incubated with 25μg Strep tag-fusion proteins (scFvs 4-2 and 22-1) and left rotating overnight at 4°C overnight. Following overnight rotation, then 40μl of Strep 2 Mag beads (IBA) were added to the cell lysates and rotated at 4°C for 1 hour. The samples were then centrifuged at 3,000 rpm for 1 minute at 4°C and supernatant discarded. Samples were then washed four times in 1 mL of Buffer W buffer, centrifuging at 3,000 rpm at 4°C for 1 minute and discarding the supernatant each time. 50 μL of Laemmli sample buffer was added to the samples, heated to 85°C for 2 minutes, centrifuged at 13,000 rpm for 10 minutes and resolved by SDS-PAGE and Immunoblot as described above. The membranes were blocked with 5% non-fat dry milk. Membranes (immunoprecipitated complexes) were probed with the mouse (anti-Zaire ebola antibody 6D8, anti-Sudan ebola virus antibody, and anti-His antibody) and rabbit (anti-Reston GP polyclonal antibody, anti-Bundibugyo GP polyclonal antibody, anti-MARV GP polyclonal antibody, and anti-Tai Forest virus GP IgG) as primary antibodies at a dilution of 1:5000 and the HRP-conjugated donkey anti-mouse and goat anti-rabbit as secondary antibodies at a dilution of 1:3000 diluted in TBST was added for a 1-hour incubation.

#### Immunofluorscence assay (IFA)

The EBOV, SUDV, BDBV, RESTV, and TAFV viruses were inoculated into 293T cells seeded in eight wells chamber slides (BD Biosciences, San Jose, CA). Forty-eight hours post inoculation infected cells were fixed with 4% paraformaldehyde solution for 10 minutes at room temperature. After fixation slides were washed three times with PBS, three minutes per wash, and blocked with PBS-0.1% BSA. Then cells were washed and incubated for one hour at room temperature with a 1:100 dilution of the ZEBOVGP scFvs 4-2 and 22-1 (500 μg /mL). After incubation, cells were washed three times with PBS, three minutes per wash. Cells were incubated at room temperature with a 1:200 dilution of goat anti-Mouse IgG Secondary Antibody, Alexa Fluor 488 (ThermoFisher) diluted in blocking buffer (PBS with 0.1% of BSA) for 1 hour at room temperature. Cells were washed three times as described above, air dry, and covered with DABCO polyvinvl alcohol mounting medium (SIGMA -ALDRICH, St. Louis, MO), followed by fluorescent microscope examination.

## Supporting information

**S1 Fig.**
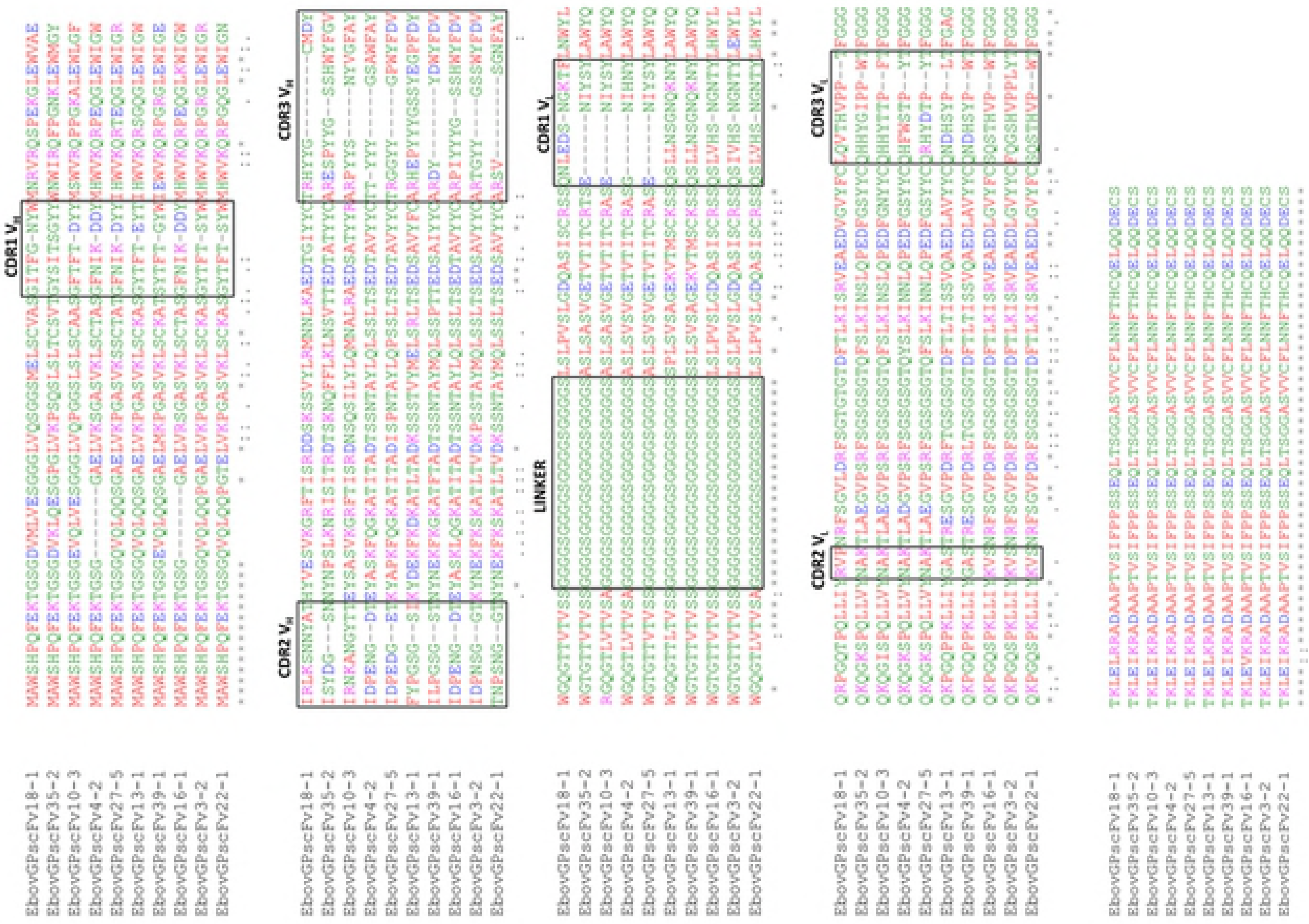
Alignment of the derived aminoacids sequences of the randomly chosen scFvs from linked antibody library and complementary determining regions (CDRs). FRs and CDRs are determined by the IMGT information system. Diversity was found predominantly in the CDR regions. A normal 20 amino acid linker [(G4S)4] joins the VH and VL chains. Alignments were **colour coded** according to residue property groups. AVFPMILW-red, DE-blue, RK-magenta, STYHCNGQ-green, others-grey. Symbols in the alignment are as follows: (*) indicates where there is a conserved amino acid; (:) indicates an amino acid position with conserved similarity; (.) indicates a semi-conserved amino substitution has occurred; (-) indicates spaces introduced to optimise the alignment.

**S2 Fig.**
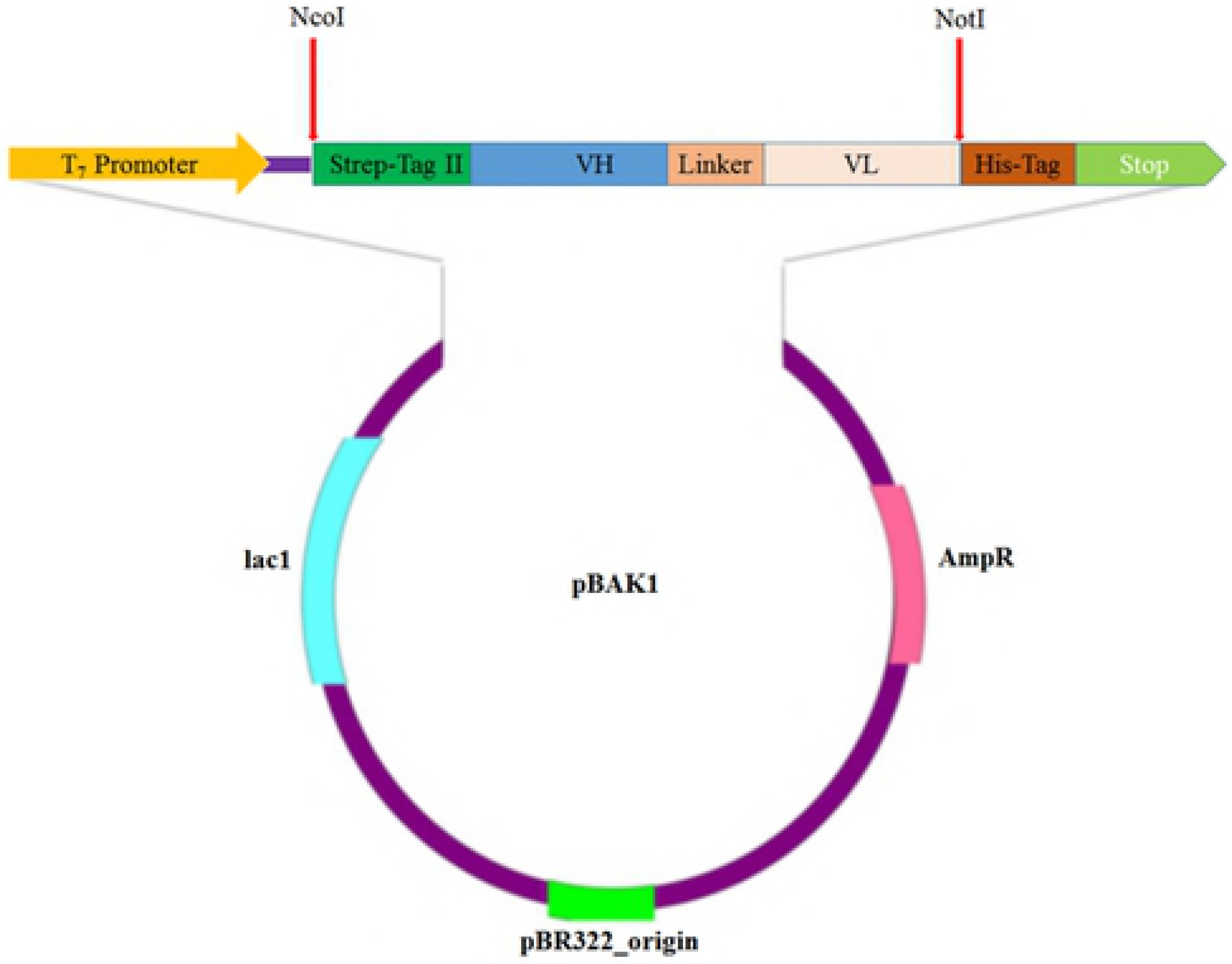
Schematic diagram of the pBAK1 (*anti-EbovGP-his-ScFv*) expression vector. The position of the enzymatic cleavage is indicated by the *red arrow*. The gene encoding anti-EbovGP-his-ScFv protein was inserted into the pBAK1 vector under the control of the T7lac promoter, in frame with a strep tag II.

**S1 Table. Nucleotide sequences of primers used.**

**S2 Table. Cross-reactivity profiles of anti-EBOV GP scFvs from ELISA result.**

**S3 Table 3. % CV & Bias around calculated EC-50.** Accuracy and precision around the calculated EC-50 values are within acceptable level.

**S4 Table. % CV & Bias around calculated EC-50.** Accuracy and precision around the calculated EC-50 values are within acceptable level.

**S5 Table. % CV & Bias around calculated EC-50.** Accuracy and precision around the calculated EC-50 values are within acceptable level.

**S6 Table. % CV & Bias around calculated EC-50.** Accuracy and precision around the calculated EC-50 values are within acceptable level.

**S7 Table. Apparent affinity and maximal binding of EBOVGP scFvs to GPZEBOV, GPSUDV, GPRESTV? GP_BDBV_, GP_TAFV_, and GP_MARV_**.

